# Circadian–Feeding misalignment in female grey mouse lemurs (*Microcebus murinus*): metabolic, physiological, and cognitive responses

**DOI:** 10.64898/2026.11.27.769665

**Authors:** Océane Lavaux, Eva Schneider, Coraline Ruiz, Jérémy Terrien

## Abstract

The consequences of eating out of phase with circadian rhythms are well documented in humans and laboratory rodents but remain poorly investigated in species that have evolved in strong seasonal constraints and fluctuating food availability. We investigated the metabolic, physiological and cognitive effects of a circadian-feeding misalignment in the grey mouse lemur (*Microcebus murinus*), a nocturnal primate characterized by pronounced metabolic flexibility. Twelve adult females underwent a Time-Restricted Feeding (TRF) protocol for 13 weeks during the summer-like season. Animals were fed either at the onset of their active phase (NIGHT-TRF, n = 6) or their rest phase (DAY-TRF, n = 6). Body composition, energy metabolism, glucose and lipid profiles, as well as learning and memory performance were monitored throughout the experiment. Feeding at the onset of the rest phase altered daily patterns of energy substrate use, whereas body mass regulation and glucose homeostasis remained largely unaffected. Although a progressive decrease in basal insulin levels as well as variations in HDL and triglyceride levels were observed, these alterations did not seem sufficient to induce a sustained metabolic syndrome. Memory performance was not affected, whereas learning abilities showed differential responses between groups. The apparent resilience of grey mouse lemurs to circadian-feeding misalignment may reflect their intrinsic metabolic flexibility, although the predictability of the feeding schedule may have facilitated gradual physiological adaptation. These findings highlight the potential of this species as a valuable model for investigating the mechanisms underlying metabolic tolerance to circadian disruption.

**Summary statement:** Feeding during the rest phase shifted the daily pattern of energy substrate use in adult female *M. murinus* without causing major deleterious effects on physiological and cognitive health, highlighting the potential role of metabolic flexibility in resilience to circadian desynchronisation.

## Introduction

Circadian rhythms are endogenous biological oscillations with a period of approximately 24 hours enabling organisms to anticipate cyclical changes in their environment. They coordinate physiological and behavioural functions, including the sleep–wake cycle, body temperature, hormone secretion, feeding behaviour, and energy metabolism (Hut & Beersma, 2011; Vitaterna et al., 2001). In mammals, the circadian system is hierarchically organised. The suprachiasmatic nuclei of the anterior hypothalamus (SCN), synchronised primarily by light–dark cycles, play the role of central clock and maintain temporal coordination throughout the organism (Longo & Panda, 2016; Mohawk et al., 2012). Peripheral clocks are also present in numerous organs involved in metabolism, including liver, pancreas, stomach, intestines, and adipose tissue. Circadian rhythms are generated by feedback loops involving genes such as *CLOCK*, *BMAL1*, *PER*, and *CRY*, while peripheral rhythms are also strongly influenced by feeding-related cues (Froy, 2007; Güldür & Otlu, 2017). Accordingly, many enzymes involved in carbohydrate, lipid, and protein metabolism, as well as hormones such as insulin, glucagon, leptin, and glucocorticoids, exhibit circadian variation. This organisation aligns nutrient absorption, storage, and utilisation with the usual feeding phase. Thus, when food is consumed outside the active phase, the light-entrained central clock may become misaligned with feeding-entrained peripheral clocks. Meal timing is therefore a major determinant of circadian coordination and metabolic health.

In humans and laboratory mice, misalignment between feeding time and the natural active phase has been associated with altered glucose and lipid metabolism, resulting in increased risk of obesity, insulin resistance, and metabolic syndrome (Froy, 2007; Güldür & Otlu, 2017; Lopez-Minguez et al., 2016). Growing evidence indicates that metabolic and circadian disturbances may also affect brain function, notably through altered sleep–wake patterns, hormone secretion, and digestive physiology disorders (Reddy & Sharma, 2026; Vaughn et al., 2014). These effects therefore extend beyond metabolism: in humans, repeated food intake outside the active phase may impair cognitive functions related to alertness, attention, memory, and information processing (Chellappa et al., 2018; Leger et al., 2018). Misalignment between endogenous rhythms and environmental cues may thus affect both energy homeostasis and cognitive performance.

Despite extensive work in humans and laboratory rodents, comparatively little is known about how feeding-induced circadian misalignment affects non-human primates, particularly species exhibiting pronounced metabolic flexibility. The grey mouse lemur (*Microcebus murinus*, Miller 1777) provides a complementary model for investigating circadian rhythms in a species exposed to pronounced environmental variation in the wild. This small nocturnal primate (50-90g body mass; 25cm long - tail included) is endemic to Madagascar, where it is exposed to substantial seasonal changes in day length, ambient temperature and food availability. As an opportunistic omnivore, it can adjust its metabolism and physiology to these changing environmental conditions (Hladik, 1979; Thorén et al., 2011). During the cold, dry season, it can enter bouts of daily torpor, which allows drastic reductions in metabolic rate and energy expenditure (Ruf & Geiser, 2015). During the warm, wet season, mouse lemurs accumulate substantial fat reserves that help meet the energetic demands of reproduction and heightened activity (Terrien, 2025). The grey mouse lemur therefore exhibits a pronounced seasonal phenotype, characterised by large changes in body mass, energy metabolism, and behaviour in both sexes, although the timing and magnitude of these changes differ between males and females (Terrien, 2025). Its capacity to transition between metabolic states reflects considerable metabolic flexibility shaped by its life history (Giroud et al., 2008; Terrien, 2025; Terrien et al., 2009), which is retained in captivity. Whether this remarkable metabolic flexibility increases resilience to feeding-induced circadian disruption or conversely, imposes additional physiological constraints, remains to date unknown.

In this study, adult female grey mouse lemurs were subjected to a Time-Restricted Feeding (TRF) protocol in which access to food was limited to a defined daily window. The control group (NIGHT-TRF) was fed at the beginning of the night, corresponding to the active phase and usual feeding period of this nocturnal species. The desynchronised group (DAY-TRF) was fed at the beginning of the day, corresponding to the usual resting phase. We aimed to characterise the response of grey mouse lemurs to feeding-induced circadian misalignment by assessing (1) physiology, particularly changes in body mass and body composition; (2) energy metabolism and the timing of carbohydrate and lipid utilisation; (3) regulation of blood glucose and circulating lipids; (4) cognition by assessing learning and memory abilities using a visual discrimination task. Given the pronounced metabolic flexibility of this species, we predicted that physiological, metabolic and cognitive responses to feeding-induced circadian misalignment would differ from those typically reported in humans and laboratory rodents, with probably less deleterious effects on the grey mouse lemurs’ health.

## Material and Methods

### Animals and ethical considerations

Twelve female grey mouse lemurs (*Microcebus murinus*), aged 2–6 years and born in the breeding colony of Brunoy (MNHN, France, licence approval no. G91-114-1) were included in the experiment. Animals were tested over a 13-week period under summer-like photoperiodic conditions (14 hours light / 10 hours dark per day). All experimental procedures were approved by the animal welfare council of UMR 7179, the Cuvier Ethics Committee for Animal Experimentation, and authorised by the French Ministry of Higher Education, Research and Innovation (authorisation no. APAFIS #59509-2026021918277341 v3). This project therefore complied with European ethical regulations governing the use of animals in biomedical research. Animals were housed in single-sex social groups of 2 to 6 individuals in aviaries, under the standard husbandry conditions of the breeding colony. Enclosures were equipped with wooden nesting boxes designed to mimic natural sleeping sites of grey mouse lemurs, such as tree hollows and cavities (Lutermann et al., 2010; Radespiel, 2006), and were enriched with branches and various objects to promote natural behavioural and physiological traits. Ambient temperature in the animal houses was maintained between 24 and 26°C and relative humidity averaged 55%.

### Project timeline

The animals were followed over a 13-week period during the summer-like season. The first three weeks, designated “W0¹”, “W0²”, “W0³” (Fig. 1), constituted a baseline phase during which all parameters were measured while all animals exhibited normal circadian and feeding schedule. The circadian-feeding desynchronisation protocol began in Week 1 (W1) in the DAY-TRF group (Fig. 1); in contrast, animals in the control NIGHT-TRF group remained fed on a normal nocturnal schedule till the end of the study.

**Figure 1:**
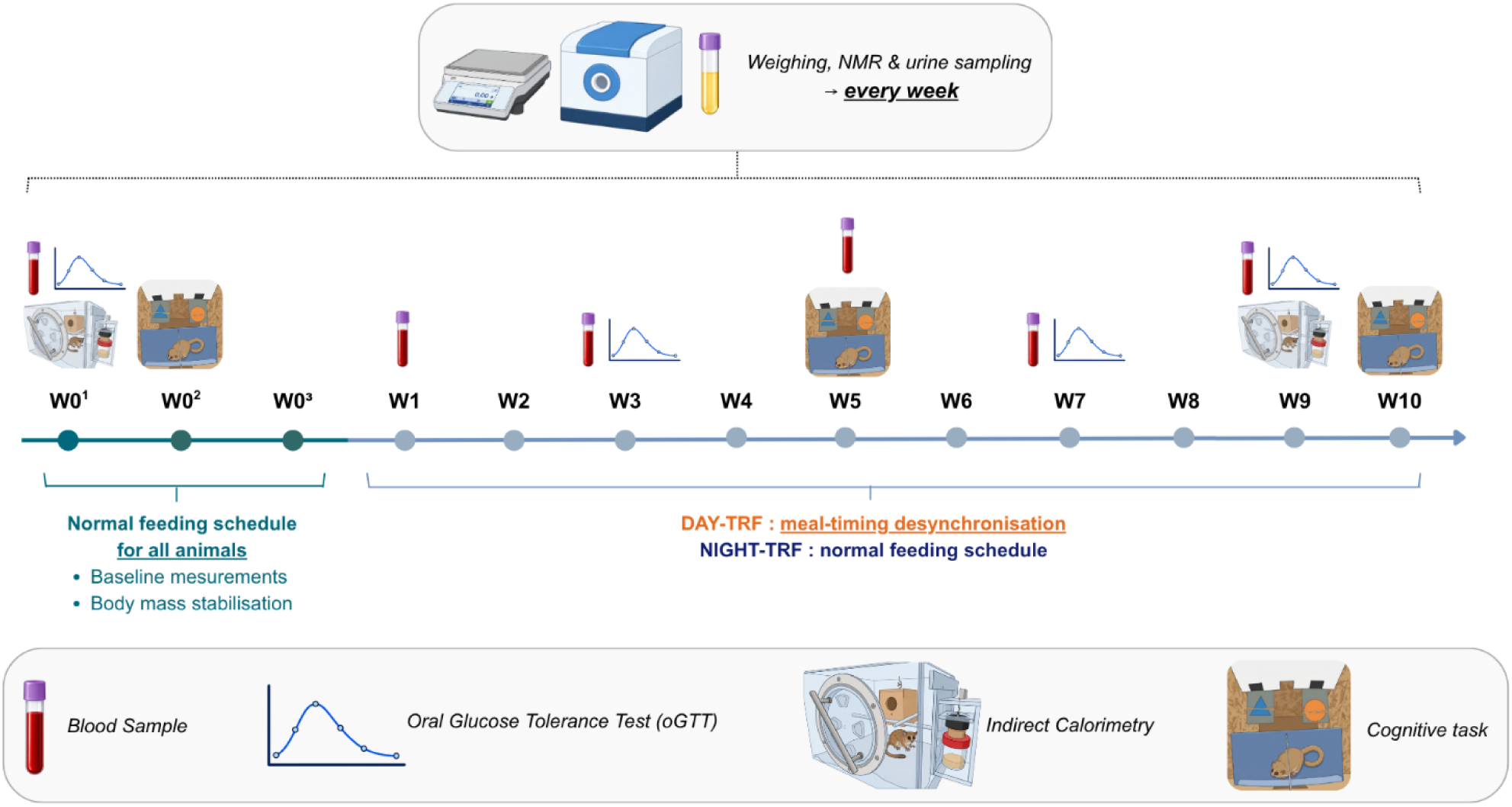
Illustration of the timeline of the Circadian–Feeding misalignment study in adult female grey mouse lemurs. The various assessments, including weighing, Nuclear Magnetic Resonance (NMR), urine and blood sampling, oral Glucose Tolerance Test (oGTT), indirect calorimetry and cognitive testing (visual discrimination task) were conducted over a 13-week period in the both control (NIGHT-TRF) and desynchronised (DAY-TRF) groups. Study weeks are designated by the letter “W”; for example, W1 corresponds to the first week of diurnal feeding in the DAY-TRF group. All animals remained under a normal feeding schedule during W0^1^, W0^2^ and W0^3^, which were used to collect baseline measurements for all parameters and allow body mass stabilisation in both groups. Illustrations were created using BioRender AI (Biorender.com) to stylize original photographs taken by the authors; the final figure layout was created manually using Canva.com.

### Circadian–Feeding misalignment protocol

The control group (NIGHT-TRF) was given daily access to food from 5:00 PM, when the lights in the animals’ facility were switched off, corresponding to the onset of the active phase and the normal feeding period for these nocturnal animals. In contrast, the desynchronised group (DAY-TRF) received food daily from 3:00 AM, when the lights were switched on, using programmable food dispensers equipped with ice cubes (VKIMISS F19-L Automatic Pet Feeder). Animals were fed a homemade mixture consisting of water, gingerbread, cereals (Blédine), powdered milk, yoghurt (8% fat), sour cream (30% fat), egg yolk and rapeseed oil. This diet provided 60% of energy coming from carbohydrates, 20% from proteins and 20% from fat, with a caloric density of 1.3 kcal/g. The same recipe was used for both NIGHT-TRF and DAY-TRF groups.

As several of the measured parameters are influenced by body mass and physical condition, it was essential to ensure that animals in the two groups had comparable body mass at the start of the experiment. Before the beginning of the study, NIGHT-TRF animals, were considerably heavier than the DAY-TRF animals (by +21% on average, and up to +40% for some individuals), and were therefore put on a restricted diet for several weeks prior the beginning of monitoring. From W0^1^, these animals received 15g of food per day, equivalent to 19.5 kcal/animal/day. To match the feeding regimen of the NIGHT-TRF group, DAY-TRF animals were initially provided with 15 g/animal/day during W0^1^ and W0^2^. However, because they lost an average of 5.5g over these two weeks (with some individuals losing up to 10g within only 10 days), the daily ration was increased to 20g per day, corresponding to 26 kcal/animal/day, at the end of W0^2^. These respective feeding regimens enabled animals in both groups to maintain a stable body mass of around 80g at the end of the baseline period, which is considered normal for the summer-like season (Fig. 3A). Both groups remained on these respective daily rations until the end of the study.

### Monitoring of body mass and composition and assessment of energy metabolism

Body mass and body composition, were monitored weekly. Animals were weighed using a precision balance (Mettler Toledo GmbH scales, model ML3002T/M00) and then inserted into an NMR analyser (Nuclear Magnetic Resonance – Mini Spec LF90II, Bruker) to accurately determine fat mass (FM) and lean body mass (LBM). The proportion of each component was then calculated and expressed as a percentage of total body mass.

Indirect calorimetry measurements were performed in W0^1^ and W9 (Figure 1) using an Oxymax system (Columbus Instruments Inc), consisting of six individual 35-liter airtight and insulated Plexiglas cages equipped with a nesting box and a feeder, and continuously supplied with fresh air at a constant flow rate of 3 litres per minute. This system enabled continuous measurement of O_2_ consumption (VO_2_, ml/kg/h) and CO_2_ production (VCO_2_, ml/kg/h), with one measurement recorded every 17.5 minutes per animal over a period of 4 to 5 days. The two first days were considered an acclimation period to the apparatus, whilst the remaining days were used for data analysis. VO_2_ and VCO_2_ were calculated using Lighton’s equations, which incorporate the LBM of each animal, previously determined by NMR (Lighton, 2008). The Respiratory Exchange Ratio (RER = VCO_2_ / VO_2_) was used to determine the nature of the metabolic substrate oxidised by the animal, following Lusk’s oxidation table (Lusk, 1924). An RER of 0.7 indicates exclusive use of lipids and is characteristic of a fasting-like metabolic state. Conversely, an RER of 1 indicates that carbohydrate oxidation is the sole source of metabolic energy. RER values greater than 1 reflect de novo lipogenesis, whereby glucose is converted into lipids for storage (neolipogenesis) (Noiret et al., 2020). Intermediate RER values around 0.8 are indicative of the simultaneous oxidation of both lipids and carbohydrates (Lighton, 2008). Daily energy expenditure (DEE) was also calculated following the equation from Lusk’s (1924) table: DEE = ((3,815 + 1,232 × RER) × VO_2_ / 1000 × LBM / 1000) (Bruss et al., 2010; Nie et al., 2015; Weir, 1949). Using this data, we also calculated the daily energy balance (DEB) over a given 24-hour period = daily caloric intake from food – daily energy expenditure (DEE).

### Assessment of Blood Glucose and Lipid Homeostasis

Blood samples were collected every two weeks, without anaesthesia. The fur on the posterior surface of the left hind limb was shaved to expose the saphenous vein, and a puncture was made using a sterile lancet. Blood was collected into EDTA-coated microcapillaries (0.1 M; 1 to 4 capillaries, corresponding to a maximum blood volume of 200 µL per animal). Samples were then centrifuged at 4500 rpm for 30 minutes at 4°C. Following centrifugation, plasma was isolated and stored at –20°C until analysis. At each sampling time point, fasting blood glucose concentration (mg/dL) was measured using a handheld glucometer (OneTouch Verio Reflect®, LifeScan). Circulating lipids profiles were also assessed using a handled lipid analyser (Mission® Cholesterol Meter, ACON LABS INC) to measure total cholesterol (CHOL), high-density lipoproteins (HDL) and triglycerides concentrations (mg/dL). Low-density lipoprotein (LDL) levels (mg/dL) were calculated using the following formula: LDL = Total cholesterol – HDL – (Triglycerides/5) (Friedewald et al., 1972).

Every second blood sampling session, i.e. every 4 weeks (Figure 1), blood sample was followed by an oral glucose tolerance test (oGTT). Briefly, animals received an oral glucose load (1.75 g of glucose per kg of body mass) and blood glucose concentration (mg/dL) was monitored 15, 30, 60 and 90 minutes after administration (Djelti et al., 2016). Based on the blood glucose concentration kinetics, glucose tolerance was defined by calculating the area under the curve (AUC, mg.dL.min^-1^) for each response using the trapezoidal method and the dplyr (1.1.4) and tidyr (1.3.2) packages in RStudio. Plasma collected on oGTT days (before glucose administration) was used to measure basal insulin concentrations (μU/mL) by ELISA (Human Insulin ELISA KIT, Millipore®, EZHI-14K) while plasma collected during routine blood sampling sessions was used to measure thyroxine (nmol/L) (T_4_ ELISA kit, LDN®, TF E-2400) in accordance with the manufacturers’ recommendations. Where insulin concentration below the assay’s detection limit, the limit of detection (LOD) was assigned and calculated based on the measured optical density (OD) of the blanks: 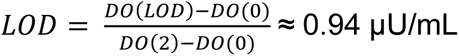 (then multiplied by 2 or 4 depending on the sample’s dilution factor). Insulin resistance was calculated using the HOMA-IR index (Homeostasis Model Assessment for Insulin Resistance) as follows: HOMA-IR = Fasting insulin (µU/mL) * Fasting blood glucose (mmol/L) / 22.5 (Marchal et al., 2012).

### Learning and memory abilities

All animals underwent a visual discrimination task under control conditions (W0² – normal night-time feeding), to establish a baseline condition unaffected by the treatment, and were subsequently tested after 5 and 10 weeks of treatment. We applied the same method as described previously (Mortessagne et al., 2024; Royo et al., 2018), using an apparatus built in Brunoy. The visual discrimination test is based on an associative learning task adapted to arboreal behaviour of the grey mouse lemur. Briefly, after a habituation phase, the animal must jump from a starting platform onto one of two platforms associated with distinct visual stimuli differing in shape, texture and colour. The correct platform provides access to the nest box, which serves as a reward, whereas the incorrect platform tips over under the animal’s weight, causing it to fall onto a protective cushion. The position of the correct platform varies randomly between trials to minimise lateral bias. The animal must therefore learn to discriminate the stimulus associated with the reward (access to the nest box); its performance being assessed based on the proportion of correct choices. The learning phase consists of a maximum of 30 trials. Each trial lasts a maximum of 5 minutes. The animal must meet a learning criterion set at 8 successful trials out of 10 consecutive ones in order to validate the test and bring it to an end. If the animal does not meet the success criterion within the 30 or ‘refuses’ the test on 5 consecutive trials, the test ends and is repeated the following day. The memory phase involves reusing the same pair of stimuli as those used during the last learning session 5 weeks earlier to check whether the animal remembers the symbol representing the positive platform. This test also consists of a session of up to 30 trials, during which the animal must meet the criterion of 8 successful trials out of 10 consecutive attempts. The parameters recorded are the total number of trials, the total number of successful trials, the total number of failures and the number of refusals. The proportion of each of these parameters is expressed as a percentage of the total number of tests.

### Urinary hormone assays

Urine samples collected weekly (Fig.1) at roughly the same time (between 1:00 PM and 2:00 PM) and on the same day (Thursday or Friday) were used to measure cortisol (Cortisol Urine ELISA, LDN, E-5100) and melatonin sulphate (Melatonin-Sulfate Urine ELISA, IBL International GmbH, RE54031) concentrations. The values for each of these parameters were corrected for urinary creatinine using the same samples (MicroVue™ Creatinine Assay Kit, Quidel Corporation, 8009). Each kit was used in accordance with the manufacturers’ recommendations. Urinary cortisol corrected for creatinine was calculated using the following formula: cortisol (ng/mg of creatinine) = cortisol (ng/mL) / (creatinine (nmol/L) * 0.11364). The same formula and units were used for melatonin sulphate values (ng/mg of creatinine).

### Statistical analyses

Statistical analyses were performed using RStudio 4.5.2 (version 2025.10.31, Posit Software, PBC). Parameters were analysed using linear mixed-effects models to account for the presence of repeated measurements within individuals, with the nlme (3.1-168) and lmerTest (3.2-1) packages. Where the assumptions of normality of residuals and homoscedasticity were met (verified using Shapiro-Wilk tests and diagnostic plots), an analysis of variance (ANOVA) and, where appropriate, post-hoc tests were applied using the emmeans (2.0.1) package. P-values for post-hoc comparisons were adjusted for multiple testing using the Holm method. Blood lipid levels, fasting blood glucose, HOMA-IR, AUC, insulin levels, urinary cortisol and melatonin sulphate concentrations, as well as success rates for both learning and memory phases of the cognitive task, were log-transformed to correct models prior to analysis. Where the conditions for application were not met, the non-parametric Wilcoxon test was used. The graphs were also created using RStudio and packages ggplot2 (4.0.1), dplyr (1.1.4) and tidyr (1.3.2) when needed. A principal component analysis (PCA) was also carried out to explore the relationships between metabolic variables (RER, 24h Energy Expenditure, T_4_, insulin, AUC, fasting blood glucose, triglycerides, cholesterol, HDL), physiological variables (body mass, urinary cortisol, urinary melatonin sulphate) and cognitive variables (learning, memory), and to visualise how individuals were categorised according to their experimental group. The variables were centred and scaled prior to analysis to ensure they were given equal weight. PCA was performed using the FactoMineR (2.14) package, and the graphical representations were produced using the factoextra (2.0.0) package. Individual coordinates on the principal dimensions were then compared between groups using a Wilcoxon test. For all analyses, we considered a difference to be statistically significant when the p-value (hereafter, p) was lower than 0.05. The data that support the findings of this study and the R code are available from the corresponding author upon reasonable request.

## Results

### Energy metabolism and body composition

Indirect calorimetry data (VO_2_, VCO_2_, RER, energy expenditure (EE)) could not be recorded for one individual in the DAY-TRF group during the week 9 session because it had to be removed from the apparatus (the animal was too stressed and did not feed). The remaining animals consumed all the food provided each day they spent in the apparatus (NIGHT-TRF: 19.5 kcal/day/animal; DAY-TRF: 26 kcal/day/animal).

In NIGHT-TRF control animals, the nocturnal phase was characterised by high mean values of VO_2_ (4227 ± 333 ml/kg/h), VCO_2_ (3625 ± 295 ml/kg/h), RER (0.86 ± 0.01) and energy expenditure (1.05 ± 0.08 kcal/h), reflecting a high level of activity during the active period (Table 1, Fig.2A and 2B, Fig.S1A and S1B). These parameters decreased significantly during the daytime phase, which corresponds to the resting period (VO_2_: 2688 ± 315 ml/kg/h; VCO_2_: 1904 ± 229 ml/kg/h; RER: 0.71 ± 0.01; DE: 0.64 ± 0.06 kcal/h) (Table 1). In animals exposed to circadian desynchronisation (DAY-TRF group), values during the nocturnal phase were also elevated, but tended to be lower than those observed in the NIGHT-TRF group for VO2 (4056 ± 280 ml/kg/h), VCO2 (3114 ± 209 ml/kg/h) and energy expenditure (0.99 ± 0.08 kcal/h) (Table 1, Fig.2B, Fig. S1A and S1B); these differences were not statistically significant between the groups (Table 1). During the daytime phase, desynchronised animals exhibited slightly higher values of VO_2_ (2886 ± 210 ml/kg/h), VCO_2_ (2531 ± 194 ml/kg/h) and energy expenditure (0.73 ± 0.08 kcal/h) than NIGHT-TRF animals; however, none of these differences reached statistical significance (Table 1, Fig.2B, Fig. S1A and S1B).

**Figure 2:**
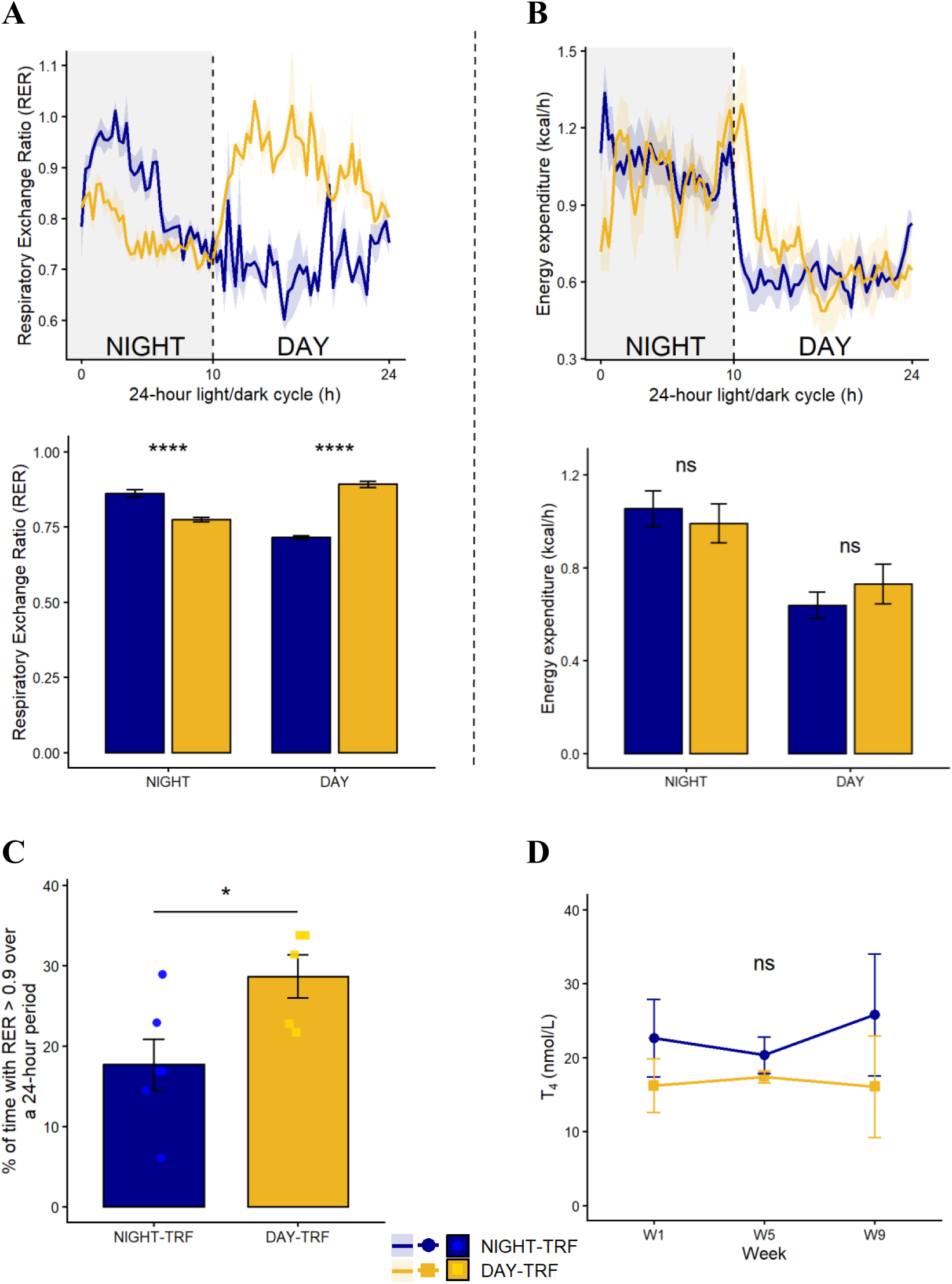
Results of the indirect calorimetry assessment performed at week 9 of the study in adult female grey mouse lemurs, subjected or not to feeding-induced circadian desynchronisation. A/ Respiratory exchange ratio (RER): mean daily profiles (top) and averaged values (bottom) according to light phase (DAY VS NIGHT) and feeding schedule (NIGHT-TRF: n = 6; DAY-TRF: n = 5) after 9 weeks of treatment. Statistical results correspond to post-hoc comparisons between light phases and feeding schedule. B/ Energy expenditure (kcal/h): mean daily profiles (top) and averaged values (bottom) according to light phase (DAY VS NIGHT) and feeding schedule (NIGHT-TRF: n = 6; DAY-TRF: n = 5) after 9 weeks of treatment. Statistical results correspond to post-hoc comparisons between light phases and feeding schedule. C/ Percentage of time spent with an RER > 0.9 over a 24-hour period, according to feeding schedule (NIGHT-TRF: n = 6; DAY-TRF: n = 6; exact Wilcoxon-Mann-Whitney Test; p=0.04). D/ Plasma thyroxine (T_4_) concentration (nmol/L) measured throughout the study according to feeding schedule (NIGHT-TRF: n = 6; DAY-TRF: n = 6). No significant effect of feeding schedule was detected (linear mixed-effects model; p = 0.2).

**Table 1:** Mean values (± SEM) of VO_2_ (ml/kg/h), VCO_2_ (ml/kg/h), RER and energy expenditure (kcal/h) measured in adult female grey mouse lemurs 9 weeks after the start of the experiment according to feeding schedule (NIGHT-TRF: n=6; DAY-TRF: n=5) and light phase (Day VS Night). Results of the statistical analyses are also presented including ANOVA table for each variable with feeding schedule (NIGHT-TRF vs DAY-TRF), light phase (Day VS Night) and their interaction. Statistical analyses were performed on individual daytime and night-time means using linear mixed-effects models, with feeding schedule, light phase, and their interaction included as fixed effects and individual identity as a random intercept.

| ***Variable*** | **VO2 (ml/kg/h)** |  | **VCO2 (ml/kg/h)** |  | **RER** |  | **Energy expenditure (kcal/h)** |  |
|---|---|---|---|---|---|---|---|---|
| *Light*<br>*Group* | Day | Night | Day | Night | Day | Night | Day | Night |
| **NIGHT-TRF** | 2688 ± 315 | 4227 ± 333 | 1904 ± 229 | 3625 ± 295 | 0.71 ± 0.01 | 0.86 ± 0.01 | 0.64 ± 0.06 | 1.05 ± 0.08 |
| **DAY-TRF** | 2886 ± 210 | 4056 ± 280 | 2531 ± 194 | 3114 ± 209 | 0.89 ± 0.01 | 0.77 ± 0.01 | 0.73 ± 0.08 | 0.99 ± 0.08 |
| ***Statistics*** | **F** | ***p*** | **F** | ***p*** | **F** | ***p*** | **F** | ***p*** |
| Feeding schedule | 0.001 | 0.97 | 0.04 | 0.86 | 21.57 | 0.0001 | 0.02 | 0.88 |
| Light | 57.98 | <0.0001 | 59.43 | <0.0001 | 7.22 | 0.02 | 52.90 | <0.0001 |
| Feeding schedule x Light Interaction | 1.04 | 0.3346 | 13.16 | 0.006 | 184.75 | <0.0001 | 2.64 | 0.14 |

The most pronounced effect of desynchronisation was observed for the respiratory exchange ratio (RER) (F = 21.57; p = 0.0001), as clearly illustrated by the daily profile in Figure 2A. At night, animals fed during the day (DAY-TRF) exhibited a significantly lower mean RER (0.77 ± 0.01) than that of the NIGHT-TRF group (0.86 ± 0.01) (p = 0.0001). By contrast, during the daytime phase, the DAY-TRF group displayed a mean RER of 0.89 ± 0.01, which was significantly higher than that of the NIGHT-TRF group (0.71 ± 0.01) (p < 0.0001) (Fig. 2A, Table 1). Furthermore, over a 24-hour period, animals subjected to DAY-TRF spent 29 ± 3% of the time with an RER > 0.9, compared with 18 ± 3% for NIGHT-TRF animals. This corresponds to a 1.5-fold increase and indicates a significantly prolonged utilisation of carbohydrate-derived substrates in DAY-TRF animals (Wilcoxon; p = 0.04) (Fig. 2C). Plasma T_4_ levels did not differ significantly between the NIGHT-TRF and DAY-TRF groups (F = 1.69; p = 0.2), and remained stable across the study (F = 0.17; p = 0.8). Over the three weeks of measurement, both groups exhibited comparable T_4_ concentrations, averaging 20 ± 5 nmol/L (Fig. 2D).

As shown in Figure 3A, the temporal reorganisation of energy substrate utilisation did not result in any body mass gain in desynchronised animals (DAY-TRF) throughout the study. Mean body mass did not differ significantly between groups (F = 0.017; p = 0.9) or over the weeks (F = 0.95; p = 0.5), with an overall mean body mass of 79 ± 1g across all individuals. Similarly, neither fat mass percentage (F = 0.024; p = 0.9) nor lean mass percentage (F = 0.99; p = 0.3) differed significantly between groups throughout the study (see for example body composition at week 9 in Fig. 3B). Furthermore, Figure 3C shows that average daily energy expenditure at week 9 did not differ significantly between groups, with animals expending approximately 20 kcal/day on average (F = 0.05; p = 0.8). However, indirect calorimetry measurements obtained during week 9 revealed that the DAY-TRF group exhibited a daily energy balance of 5.9 ± 1.8 kcal/day, which was significantly higher than that of the NIGHT-TRF group (−0.4 ± 1.4 kcal/day) (F = 6.88; p = 0.03) (Fig. 3D). Hence, despite a higher positive daily energy balance as revealed by indirect calorimetry, desynchronised animals showed no evidence of obesity or fat mass gain, or any major thyroid dysfunction throughout the study.

**Figure 3:**
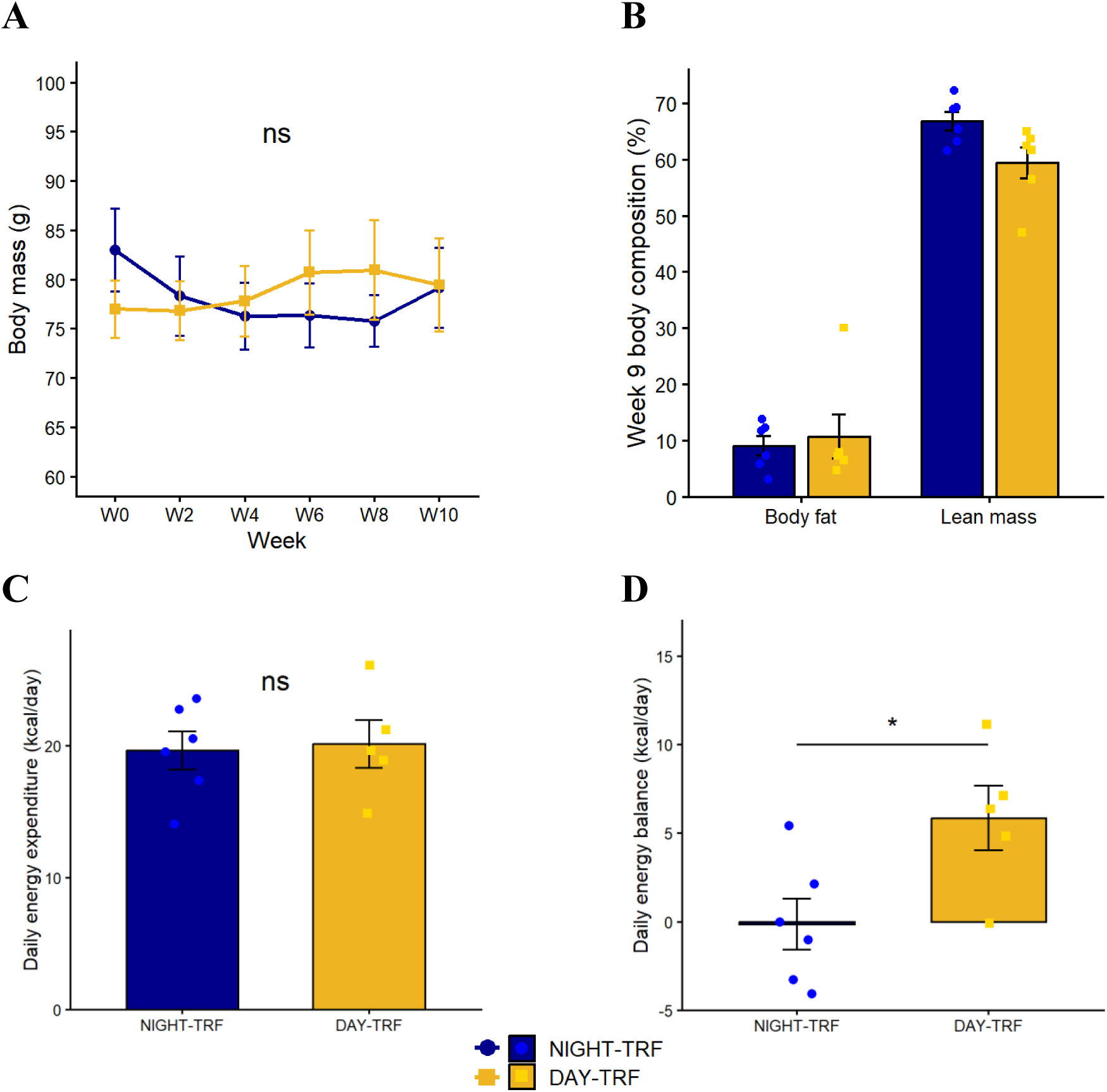
Effects of feeding-induced circadian desynchronisation on body composition, daily energy expenditure and energy balance in adult female grey mouse lemurs. A/ Changes in mean body mass (g) throughout the study according to feeding schedule (NIGHT-TRF: n = 6; DAY-TRF: n = 6). No significant effect of feeding schedule was detected (p = 0.9). B/ Mean body composition (% of fat mass and % of lean mass) measured during week 9 in both groups (NIGHT-TRF: n = 6; DAY-TRF: n = 6). C/ Mean daily energy expenditure (kcal/day) measured during week 9, according to feeding schedule (NIGHT-TRF: n = 6; DAY-TRF: n = 5). No significant effect of feeding schedule was detected (linear mixed-effects model; p = 0.8). D/ Mean daily energy balance (kcal/day) measured during week 9, according to feeding schedule (NIGHT-TRF: n = 6; DAY-TRF: n = 5). The DAY-TRF group exhibited a significantly higher energy balance than the NIGHT-TRF group (linear mixed-effects model; p = 0.03).

Interestingly, the temporal reorganisation of energy substrate utilisation did not appear to induce chronic stress in the desynchronised animals, as urinary cortisol levels remained stable throughout the study at around 13 ± 0.4 ng/mg of creatinine, and did not differ from those of the control group (p = 0.43) (Fig. S2A). Furthermore, synchronisation of the central clock appeared to remain robust despite the desynchronised feeding schedule, as urinary melatonin sulphate concentrations did not differ significantly between the DAY-TRF (0.35 ± 0.05 ng/mg of creatinine) and NIGHT-TRF (0.34 ± 0.07 ng/mg of creatinine) groups across the weeks measured (p = 0.42) (Fig.S2B).

### Blood lipids and glucose homeostasis

The shift in meal timing appears to have had contrasting effects on the various blood lipids parameters measured. Mean total cholesterol levels (Fig. 4A) did not differ significantly between the desynchronised group (DAY-TRF: 262 ± 27 mg/dL) and the control group (NIGHT-TRF: 222 ± 19 mg/dL) (F = 2.5; p = 0.2) and remained relatively stable throughout the study (F = 1.9; p = 0.1). Similarly, LDL levels (Fig. 4B) showed no significant variations over time (F = 2.2; p = 0.1), averaging 150 ± 26 mg/dL in the DAY-TRF group and 111 ± 19 mg/dL in the NIGHT-TRF group, with no significant difference (F = 0.04; p = 0.9). In contrast, desynchronised animals exhibited mean HDL levels (Fig. 4C) of 97 ± 2 mg/dL which were significantly lower (W = 55; p = 0.001) than those of the control animals, all of which remained at the assay’s upper detection limit of 100 mg/dL throughout the study. However, this difference was no longer evident by the ninth week (p_W9_ = 0.2).

**Figure 4:**
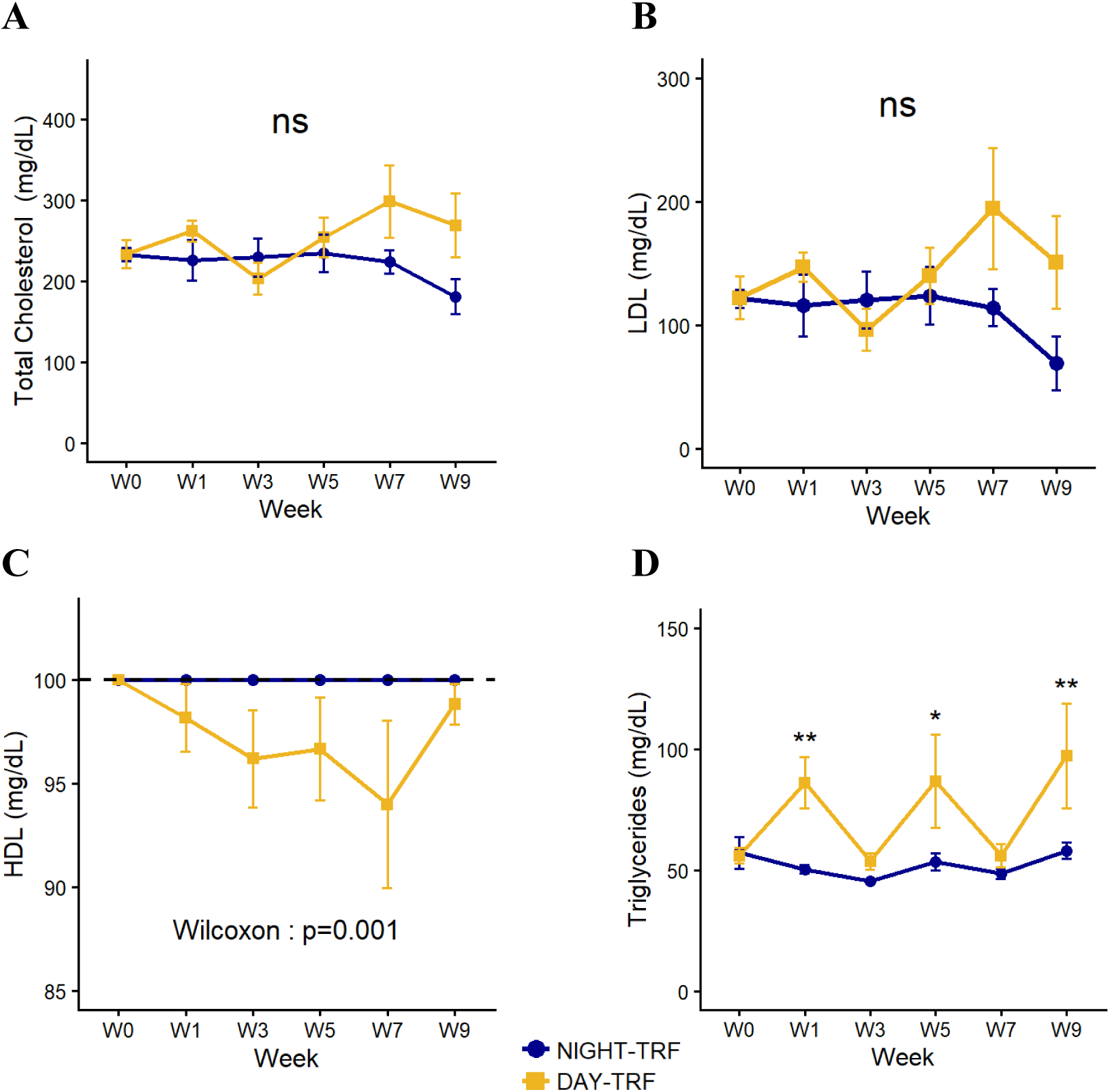
Changes in mean blood lipid concentrations (mg/dL) over time (weeks) in adult female grey mouse lemurs subjected to feeding desynchronisation (DAY-TRF: n = 6) or maintained under a synchronised feeding schedule (NIGHT-TRF: n = 6). Total cholesterol, LDL and triglyceride concentrations were log-transformed prior to analysis and evaluated using linear mixed-effects models. A/ Total cholesterol: no significant effect of feeding schedule (p = 0.2). B/ LDL: no effect of feeding schedule (p = 0.1). C/ HDL: significant effect of feeding schedule (Wilcoxon Test: p = 0.001). The dotted horizontal line indicates the analyser’s maximum detection limit. D/ Triglycerides: statistics indicate the effect of feeding schedule at each time point (post-hoc tests).

Conversely, the DAY-TRF group exhibited mean triglyceride levels (Fig. 4D) of 73 ± 10 mg/dL, which were significantly higher than those observed in the NIGHT-TRF group (52 ± 3 mg/dL) (F = 6.3; p = 0.03). Post-hoc tests revealed that the differences between groups were significant at weeks 1 (p = 0.003), 3 (p = 0.02) and 9 (p = 0.001). Overall, desynchronisation of the feeding schedule did not significantly alter total cholesterol or LDL levels but was associated with increased triglyceride concentrations and reduced HDL levels.

Plasma basal insulin measurements (Fig. 5A) showed that basal insulin levels were, on average, significantly lower in desynchronised animals than in the control group (F = 8.3; p = 0.01). Pairwise comparisons indicated no difference between the groups at week 0, when all animals are fed during the night (p = 0.6). However, whereas basal insulin levels remained relatively stable in NIGHT-TRF animals (week 3: 69 ± 26 µU/mL; week 7: 67 ± 16 µU/mL), those of the DAY-TRF group progressively decreased over time and tended to be lower than those of the NIGHT-TRF group by week 3 with a mean of 22 ± 5 µU/mL (p = 0.08). This trend became more pronounced by Week 7, with mean insulin levels reaching 7 ± 2 µU/mL, a value markedly lower than that observed in the control group (p < 0.0001). Similarly, the HOMA-IR index (Fig. S3A) gradually decreased in animals subjected to the DAY-TRF schedule, falling from 9 ± 2 at week 0 to 5 ± 1 at week 3 and then to 1 ± 1 at week 7. In contrast, HOMA-IR values remained high in the control group (12 ± 6, 13 ± 4 and 18 ± 6, at weeks 0, 3 and 7, respectively). Consequently, HOMA-IR values were significantly lower in DAY-TRF animals at both weeks 3 (p = 0.05) and 7 (p < 0.0001).

**Figure 5:**
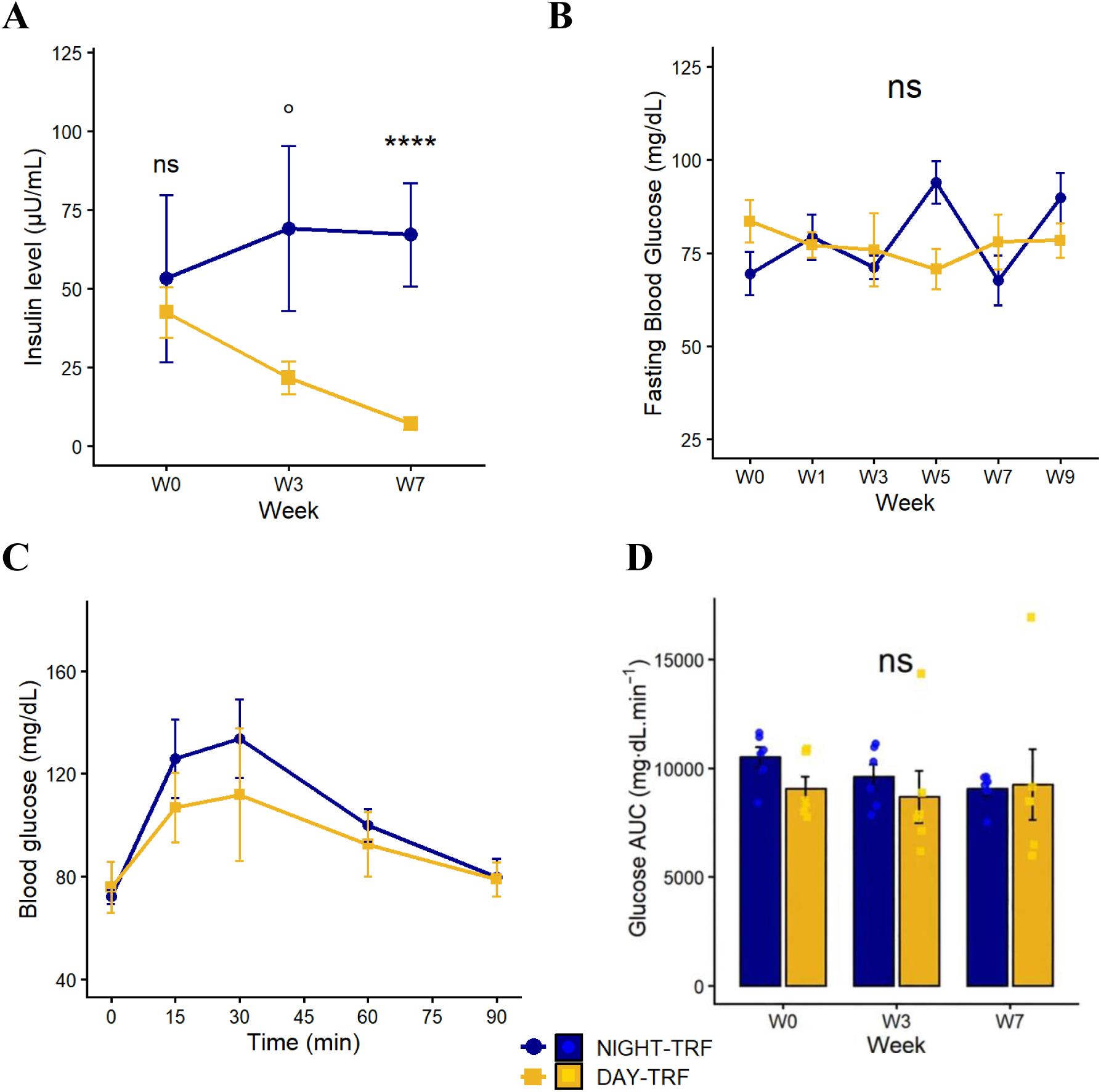
Assessment of glucose homeostasis in adult female grey mouse lemurs subjected to circadian desynchronisation (DAY-TRF: n = 6) or maintained under a synchronised feeding schedule (NIGHT-TRF: n = 6). Data were analysed using linear mixed-effects models; insulin and AUC data were log-transformed prior to analysis. A/ Changes in mean basal insulin levels (µU/mL) over the course of the study. Statistics indicate the effect of feeding schedule within each week (post-hoc tests). B/ Changes in mean fasting blood glucose levels (mg/dL) over time; no significant effect of the feeding schedule was detected (p = 0.9). C/ Mean blood glucose response curves during the oral glucose tolerance test (oGTT) performed at week 3 in both groups. D/ Mean glucose tolerance indices, expressed as the area under the curve (AUC, mg.dL.min^-1^) across the study period; no significant effect of feeding schedule was detected (p = 0.3).

Interestingly, this reduction in basal insulin levels in the DAY-TRF animals had no effect on either fasting blood glucose levels or glucose tolerance. Indeed, measurements of fasting blood glucose levels performed every two weeks revealed no significant difference between the two groups (F = 0.008; p = 0.9) (Fig. 5B). Although blood glucose levels appeared to be slightly higher in the control group at week 5, the mean values remained very similar between groups (NIGHT-TRF: 84 ± 8 mg/dL; DAY-TRF: 81 ± 8 mg/dL). At weeks 0, 3 and 7, blood sampling was followed by an oral glucose tolerance test (oGTT) to monitor blood glucose levels over a 90-minutes period following the ingestion of a glucose solution (see week 3 oGTT in Fig. 5C and week 0-7 oGTTs in Fig. S3B and S3C). Analysis of the areas under the curve (AUC) for the glucose response curves (Fig. 5D) indicated that glucose tolerance remained comparable between the two groups (F = 1.49; p = 0.3) averaging 9329 ± 60 mg.dL.min^-1^ amongst all animals. Overall, despite reduced basal insulin levels, the desynchronised animals showed no evidence of impaired glucose tolerance, insulin resistance, or diabetes.

### Cognitive consequences

The visual discrimination experiment (Fig. 6) showed that feeding-induced circadian desynchronisation had no effect on the success rate of animals in the DAY-TRF group compared with those in the control group, either during the learning phase (F = 0.009; p = 0.9) or the memory phase (F = 0.17; p = 0.8). For instance, during the learning phase, after five weeks of desynchronisation, DAY-TRF animals achieved a mean success rate of 89 ± 5%, which did not differ significantly from that of NIGHT-TRF animals (80 ± 10%). Similarly, during the memory phase, after 10 weeks of desynchronisation, the DAY-TRF group exhibited a mean success rate of 87 ± 7%, compared with 81 ± 9% in the NIGHT-TRF group, with no significant difference between groups. However, a significant interaction between feeding schedule and week was detected during the learning phase (F = 6.29; p = 0.03). Post-hoc tests revealed that the success rate of the desynchronised group was significantly lower at week 10 (69 ± 3%) than week 5 (89 ± 5%), suggesting a progressive decline in learning performance over time in animals subjected to DAY-TRF protocol (p = 0.007).

**Figure 6:**
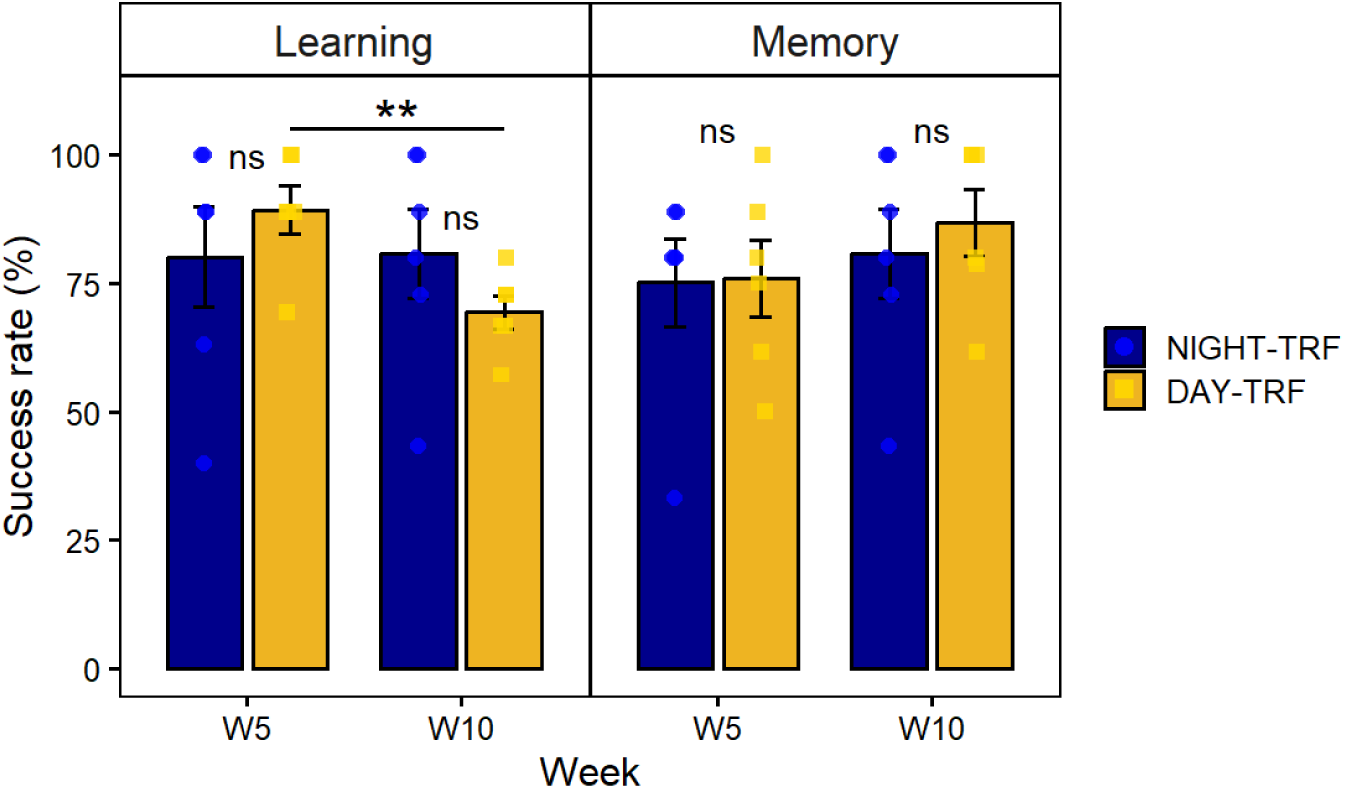
Mean success rates in the learning and memory tasks of the visual discrimination experiment in adult female mouse lemurs subjected to circadian desynchronisation (DAY-TRF: n = 6) or maintained under a synchronised feeding schedule (NIGHT-TRF: n = 6) according to study weeks. Statistics indicate the effect of feeding schedule, as well as the post-hoc comparison between the DAY-TRF group at weeks 5 and 10. Success rates for both the learning and memory phases were log-transformed prior to analysis).

### Exploratory multivariate analysis

A principal component analysis (PCA) was performed to compare the physiological, metabolic and behavioural parameters measured affected by feeding desynchronisation and to provide an integrated overview of the animals’ profiles after 10 weeks of treatment (Fig. 7). The first two principal components explained 43.8% of the total variance (Dim1: 25.1%; Dim2: 18.7%). Animals from the NIGHT-TRF and DAY-TRF groups were primarily separated along the first dimension (Wilcoxon; p = 0.004), whereas no significant separation was observed along the second dimension (Wilcoxon; p = 0.24). DAY-TRF individuals clustered on the negative side of Dim1, whereas most NIGHT-TRF animals had positive scores. Dim1 was negatively correlated with daily mean RER (r = −0.79, p = 0.002), daily energy balance (r = −0.79, p = 0.002), triglycerides (r = −0.61, p = 0.04), and total cholesterol (r = −0.6, p = 0.04), and positively correlated with insulin (r = 0.77, p = 0.004) and fasting blood glucose (r = 0.62, p = 0.03). Plasma T4 concentrations also showed a moderate positive correlation with Dim1 (r = 0.56, p = 0.060). Dim2 was positively correlated with the area under the oral glucose tolerance test curve (oGTT AUC, r = 0.8, p = 0.002), urinary melatonin sulphate (r = 0.77, p = 0.003), and fasting blood glucose (r = 0.61, p = 0.03). Body mass, urinary cortisol, daily energy expenditure, HDL as well as learning and memory abilities showed weak correlations with both axes. Overall, these exploratory analyses suggest that feeding desynchronisation induced distinct multivariate physiological and metabolic profiles. It should be noted that the reported p-values correspond to correlations between variables and principal components rather than formal tests of between-group differences.

**Figure 7:**
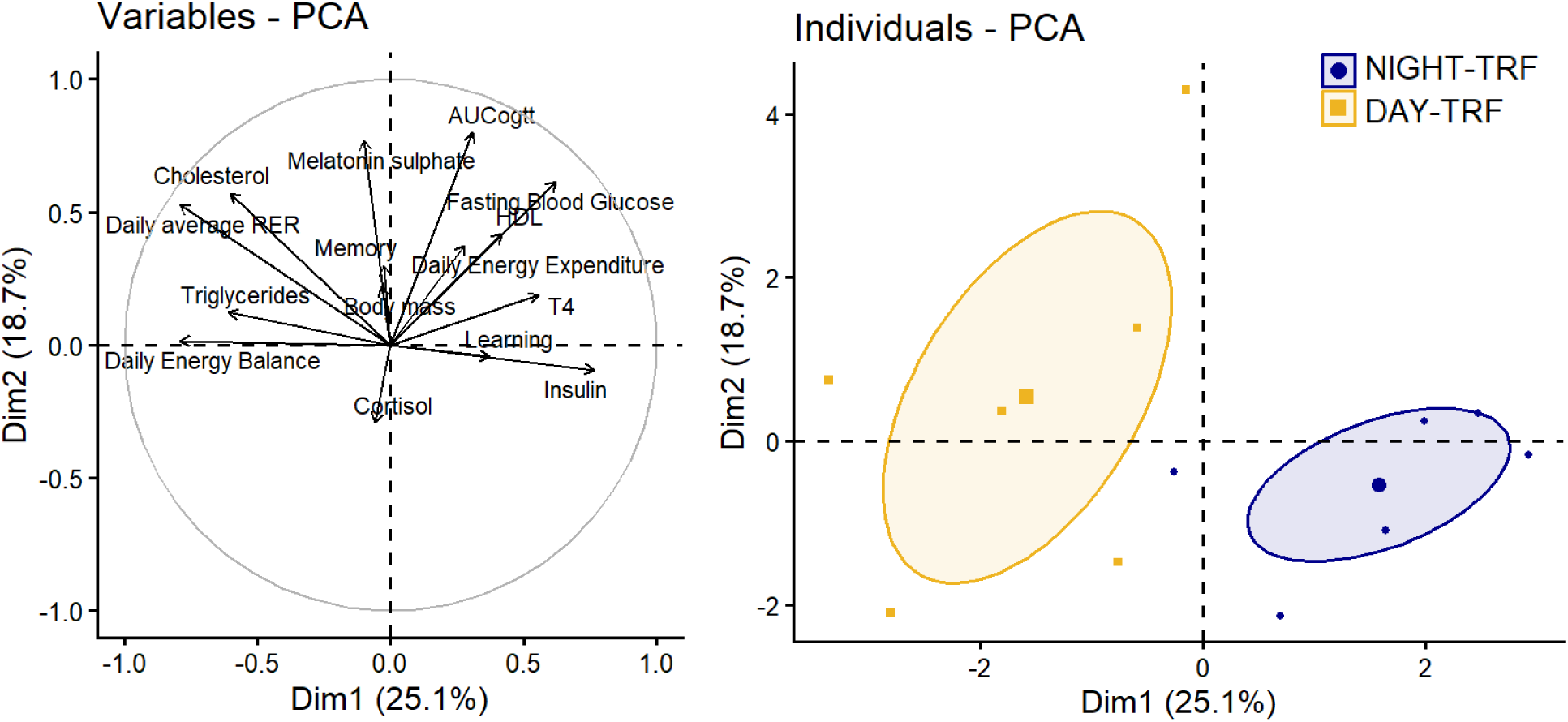
Principal component analysis (PCA) of physiological, metabolic and cognitive parameters measured in control (NIGHT-TRF: n = 6) and desynchronised (DAY-TRF: n = 6) adult female grey mouse lemurs after 8-10 weeks of treatment. The parameters included in the PCA are body mass, triglycerides, total cholesterol, HDL, RER, daily energy expenditure, plasma T_4_, glucose tolerance indices expressed as the area under the curve (AUC), fasting blood glucose, plasma insulin, urinary cortisol and melatonin sulphate, as well as success rates in memory and learning tests at weeks 8,9 or 10. The first two principal components explained 43.8% of the total variance (Dim1: 25.1%; Dim2: 18.7%). Individuals from the two groups were primarily separated along Dim1 (Wilcoxon; p = 0.004) whereas no significant separation was observed along Dim2 (Wilcoxon; p = 0.24).

## Discussion

In contrast to the detrimental effects reported in humans and laboratory rodents, circadian-feeding misalignment produced only limited physiological and metabolic alterations in female *Microcebus murinus*. The main changes observed in desynchronised animals consisted of a temporal reorganisation of energy substrate utilisation, altered triglyceride and HDL levels, and a progressive decrease in blood insulin levels, however with no effect on fasting blood glucose or glucose tolerance. Total cholesterol, LDL and plasma T_4_ levels remained unaffected. Body composition in the desynchronised animals also did not differ from that of control animals, showing no evidence of obesity. Although there may be a potential effect on learning abilities, the desynchronisation of feeding schedule had no effect on memory. This observed resilience of female mouse lemurs to adapt efficiently to a chronic shift in feeding time could then reflect the species’ exceptional innate metabolic flexibility.

### Feeding misalignment induces profound temporal reorganisation of energy metabolism

The indirect calorimetry data reported in this study are consistent with the expected physiology of the grey mouse lemur: VO_2_; VCO_2_ and energy expenditure were significantly higher at night than during the day in the control group, in accordance with the species’ nocturnal lifestyle. More interestingly, the shift in feeding schedule profoundly altered the temporal pattern of energy substrate utilisation. Indeed, analyses of the respiratory exchange ratio (RER) showed that, when feeding is aligned with the circadian rhythm, *M. murinus* relies almost exclusively on lipid reserves during the daytime resting phase (RER = 0.71 ± 0.01) and on a mixed energy source at night, predominantly derived from carbohydrates (RER = 0.86 ± 0.01). In contrast, the mean RER at night was equal to 0.77 ± 0.01 in desynchronised animals, indicating mixed lipid and carbohydrate utilisation, but more importantly, a very high mean RER of 0.89 ± 0.01 during the day, reflecting a predominant reliance on carbohydrates during the resting phase. We also found that animals fed at the onset of their rest phase spent 1.5 times longer with an RER > 0.9 than animals fed at the onset of their active phase. One hypothesis to explain this observation is that daytime feeding is misaligned with the digestive circadian clock, thereby affecting intestinal motility and nutrient absorption. Gastrointestinal motility is known to follow a circadian rhythm and is markedly reduced during the rest/sleep phase (Duboc et al., 2020). Reduced gut motility during the daytime resting phase of grey mouse lemurs may prolong carbohydrate absorption, thereby sustaining carbohydrate oxidation and explaining the extended daily duration of RER > 0.9 observed under circadian–feeding misalignment. Similar disturbances have been observed in rats fed during their resting phase, which exhibit disrupted gastrointestinal rhythms, altered gastrointestinal motility and desynchronisation of local gastrointestinal clocks (Salgado-Delgado et al., 2023). Similarly, in humans, disturbances in intestinal circadian rhythms have been associated with gastrointestinal disorders, including constipation and irritable bowel syndrome (Duboc et al., 2020). In addition to the reorganisation of energy substrate utilisation, changes in gut microbiota rhythms could also contribute to this phenomenon. Indeed, the gut microbiota exhibits robust circadian oscillations that are strongly influenced by feeding–fasting cycles: feeding during the rest phase may alter these rhythms and consequently affect microbial metabolite production, intestinal motility and nutrient absorption (Kaczmarek et al., 2017; Thaiss et al., 2014; Wu et al., 2026; Zarrinpar et al., 2014). Although these parameters were not assessed in the present study, faecal samples were collected during the weekly weighing sessions and may provide a valuable opportunity to investigate whether circadian-feeding misalignment induces changes in the gut microbiota of grey mouse lemurs.

The reversal of temporal pattern of energy substrate utilisation was not unexpected, as carbohydrates are preferentially oxidised whenever they are available: through the action of insulin, carbohydrate utilisation is promoted while lipid mobilisation and oxidation are simultaneously suppressed (Goodpaster & Sparks, 2017). In DAY-TRF animals, the normal carbohydrates-to-lipid metabolic transition was therefore shifted by the altered feeding schedule, without compromising the overall ability to switch between energy substrates. In this context, the changes in RER observed in DAY-TRF animals may reflect a very efficient adaptive reorganisation of substrate utilisation rather than a profound disruption of metabolic homeostasis. This interpretation is consistent with the absence of major changes in body mass, body composition or most of metabolic parameters, indicating that *M. murinus* possesses a remarkable capacity to maintain energy homeostasis despite a temporal constraint on food intake. Moreover, despite the pronounced reorganisation of energy metabolism induced by daytime feeding, plasma T_4_ levels remained unchanged throughout the experiment. This lack of effect is consistent with findings in both humans and rodents, where circadian desynchronisation tends to affect TSH (Thyroid-Stimulating Hormone) secretion and the temporal organisation of the hypothalamic-pituitary-thyroid axis, whereas plasma T_4_ concentrations generally remain stable due to strong homeostatic regulation and the hormone’s relatively long half-life (Jeong et al., 2023; Leso et al., 2020). Also, despite the reorganisation of the daily pattern of substrate utilisation in DAY-TRF animals, neither urinary 6-sulfatoxymelatonin nor urinary cortisol differed significantly between feeding groups at the scheduled sampling time. A similar pattern has been reported in humans, where delaying meal timing shifted the plasma glucose rhythm without affecting melatonin onset or the timing of the plasma cortisol peak (Wehrens et al., 2017). Our findings are therefore consistent with continued entrainment of the central circadian pacemaker to the light–dark cycle. However, determining whether the phase and amplitude of these hormonal rhythms were fully preserved would require sampling at multiple time points across the 24-h cycle.

### Despite modified substrate utilisation and apparent positive daily energy balance, body composition remained stable

A key limitation of the present study was the unequal daily caloric intake between groups (NIGHT-TRF: 19.5 kcal/animal/day; DAY-TRF: 26 kcal/animal/day), which was yet necessary to maintain comparable and stable body masses consistent with the summer-like physiological state of the animals. Excess body mass could have confounded metabolic outcomes, as overweight and obesity are associated with altered lipid profiles (Terrien et al., 2018), impaired glucose tolerance and insulin resistance (Ye, 2013). Moreover, increased adiposity might also have affected motivation and performance during the visual discrimination task, which required repeated jumping over sessions lasting 1 to 1.5 hours. Obesity has also been associated with cognitive impairments in both mice and humans (Cheke et al., 2016; Farr et al., 2008). The feeding regimen adopted in each group therefore helped maintain comparable physiological conditions between groups throughout the study. Interestingly, and given the known effects of feeding desynchronisation on body mass, the interaction between circadian-feeding misalignment and higher daily calorie intake in the DAY-TRF group should have led to marked obesity; however, this was not the case in *M. murinus*, reinforcing the idea that this species exhibits a high degree of resilience to environmental changes. Although, differences in caloric intake naturally remain a potential confounding factor when interpreting the daily energy balance data. Also, daily energy expenditure and energy balance were assessed during indirect calorimetry sessions, when individuals were housed separately and individual food intake could be accurately monitored. These conditions are very different from the animals’ usual housing situation, in which they share aviaries in social groups. Such housing condition may result in competition for food access and prevents accurate monitoring of individual energy intake. Consequently, social hierarchy could have influenced food consumption, with some individuals potentially consuming more or fewer calories than others throughout the study. Furthermore, because the size and environmental enrichment of the calorimetry enclosures are much more limited than in the aviaries, energy expenditure measured during indirect calorimetry sessions may underestimate that occurring under standard housing conditions, especially due to the reduced contribution of locomotor activity. Nevertheless, during the final 24 hours of the week 9 calorimetry session (W9 Day 4), under single-housing conditions, the DAY-TRF group exhibited a significantly higher mean daily energy balance (6 ± 2 kcal/day) than the control group (0 ± 1 kcal/day). A similarly positive daily energy balance was also observed on the preceding day (W9 Day 3, Fig. S4B), averaging 7.9 ± 1.4 kcal/day in DAY-TRF animals. In contrast, during the baseline session at week 0, when both groups received the same feeding schedule and caloric ration, daily energy balance remained close to the equilibrium in both NIGHT-TRF and DAY-TRF animals (Fig. S4B). These results suggest that, despite comparable daily energy expenditure across the three time points analysed in the DAY-TRF group (W0 Day 4; W9 Day 3; W9 Day 4 - Fig. S4A), the misaligned feeding schedule may have contributed to the observed alteration in daily energy balance. Logically, one might hypothesise that the positive energy balance observed in the DAY-TRF group at week 9 resulted, at least in part, from the higher daily caloric intake. However, if the DAY-TRF group (fed with 26 kcal/animal/day from the end of W0^2^) had shown a positive energy balance of +6 to +8 kcal/day throughout the study, they should have gained a significant amount of body mass and body fat. Yet, there was no evidence of obesity in desynchronised animals as the mean body mass of the DAY-TRF group remained stable throughout the study averaging 80g/individual, just like the NIGHT-TRF group. An alternative hypothesis is that feeding at an inappropriate time, particularly at the beginning of the resting phase, may alter intestinal and metabolic rhythms, thereby potentially affecting the energetic costs of nutrient processing. Importantly, the calculated daily energy balance should be interpreted as an approximation of short-term energetic status derived from indirect calorimetry measurements rather than as a direct measure of chronic energy balance. Nevertheless, analyses of faecal calorie content as well as locomotor activity during calorimetry sessions could notably allow to calculate a precise digestive cost and to support this hypothesis. Analyses of plasmatic hormones involved in digestive process could further improve our understanding of the physiological adaptations associated with circadian-feeding misalignment.

### Evidence for metabolic adaptation rather than metabolic dysfunction

At first glance, the lipid profile observed in the DAY-TRF group resembles that reported in humans and rodents subjected to circadian desynchronisation. Indeed, permanent night-shift workers typically exhibit elevated triglyceride levels, reduced HDL levels, a moderate increase in total cholesterol and little or no change in LDL concentrations (Dutheil et al., 2020; Reutrakul & Knutson, 2015). Similarly, rats fed during their resting phase show impaired clearance of triglyceride-rich lipoproteins, leading to a transient accumulation of circulating triglycerides (Opperhuizen et al., 2015). One possible explanation is that, during the resting phase, dietary lipids are processed by the liver at a time when its metabolic programmes are normally oriented toward the mobilisation of energy stores and the oxidation of fatty acids rather than lipid storage or redistribution. This mismatch between feeding time and hepatic metabolic rhythms may therefore impair triglyceride clearance, leading to elevated circulating triglyceride levels. Nevertheless, the results concerning HDL levels should be interpreted with caution. In the control group, HDL concentrations consistently reached the analyser’s upper detection limit (100 mg/dL) at every time point, resulting in zero variance. Consequently, HDL values may have been underestimated, and the absence of variance precluded the use of parametric statistical tests. However, the convergence of HDL levels between groups by week 9 of the study suggests that these differences may have been transient and that the DAY-TRF animals progressively adapted to the desynchronised feeding schedule. This interpretation is consistent with the Time-Restricted Feeding (TRF) paradigm, according to which regular feeding-fasting cycles can synchronise peripheral metabolic clocks, even when feeding occurs out of phase with the central circadian clock. Thus, the predictable daily feeding schedule may have facilitated progressive physiological adaptation in DAY-TRF animals. Specifically, TRF has been shown to prevent body mass gain, glucose intolerance and fatty liver disease in mice fed a high-fat diet, regardless of the total number of calories consumed (Regmi et al., 2021). Likewise, TRF protects against obesity and metabolic syndrome in C57BL/6J mice lacking a functional circadian clock (Chaix et al., 2019).

Desynchronised animals showed a progressive decrease in both basal insulin levels and HOMA-IR index throughout the study, however with no effect on fasting blood glucose levels or glucose tolerance. This suggests that desynchronised animals were able to maintain blood glucose levels comparable to those of control animals with significantly lower insulin levels. These findings are suggestive of improved insulin sensitivity, particularly at the hepatic level, allowing endogenous glucose production to be regulated with lower insulin levels. This metabolic profile has been reported in desynchronised rats (Pérez-Mendoza et al., 2014). It is however important to note that the HOMA-IR index was originally developed and validated in humans to assess insulin resistance. Its interpretation in grey mouse lemurs should therefore be made with caution and should primarily be used for comparative rather than diagnostic purposes. Insulin levels are also influenced by the duration of fasting following a meal. However, within the framework of the present TRF protocol, blood samples were always collected at the same time of day (1:30 PM) in both groups. Thus, given the shift in feeding schedules, there was a difference of approximately 10 hours between the groups in the duration of fasting at the time of blood sampling. Specifically, NIGHT-TRF animals were sampled after approximately 20 hours of fasting, whereas DAY-TRF animals had fasted for only 10 hours. It would therefore be interesting to collect blood samples from DAY-TRF animals at 11:00 PM to verify whether the observed insulin response is attributable, at least in part, to the difference in fasting duration between groups. Also, given this difference, one might have expected DAY-TRF animals to exhibit higher fasting blood glucose concentrations and to be closer to a post-absorptive state than NIGHT-TRF animals. Nevertheless, blood glucose levels remained comparable between groups throughout the study. A likely explanation is that, after 8 to 10 hours without food, DAY-TRF animals had already exited the postprandial phase. In other species such as mice, experimental guidelines for oral glucose tolerance test (oGTT) generally indicate that a fasting period of 4 to 6 hours is sufficient to reduce blood glucose levels and ensure gastric emptying, while a fast of 10 to 12 hours is commonly used to obtain baseline fasting blood glucose values (Benedé-Ubieto et al., 2020; Kennard et al., 2022). Similarly, in humans, fasting blood glucose levels are typically assessed after a fast of 8 to 12 hours, a duration considered sufficient to achieve metabolic steady-state conditions. For example, both glucose and insulin concentrations decrease significantly after an overnight fast (Patterson & Sears, 2017). Overall, and in contrast to findings commonly reported in humans and laboratory rodents, no evidence of insulin resistance, diabetes or sustained metabolic syndrome was observed in this study. These findings support the hypothesis that adult female *Microcebus murinus* can efficiently adapt to feeding-related circadian misalignment rather than developing metabolic dysfunctions.

### Consistency between preservation of physiological integrity and cognitive abilities

In both humans and mouse models, chronic metabolic disturbances such as insulin resistance, obesity and metabolic syndrome are frequently associated with impaired cognitive functions, particularly affecting memory and executive functions. These effects are thought to arise through multiple mechanisms, including inflammation, oxidative stress, vascular dysfunction and impaired cerebral insulin signalling (Farruggia & Small, 2019). In the present study, the overall preservation of physiological and metabolic parameters is therefore consistent with the absence of major impairments of cognitive performance in DAY -TRF animals relative to the NIGHT-TRF group. One question remains, however, regarding the final learning phase of the visual discrimination task performed during week 10. At this time point, the DAY-TRF group exhibited a significantly lower success rate than that observed during week 5, although performance did not differ significantly from that of the control group. Circadian-feeding misalignment may therefore alter learning abilities on a longer term, a hypothesis that should be investigated in studies beyond the 10-week duration of the present experiment. However, this finding should be interpreted with caution, as DAY-TRF animals displayed a clear reduction in motivation during the final day of the learning phase, an observation that was not made during the memory session conducted the previous day.

A decline in cognitive performance could also have been expected in DAY-TRF animals due to a potential reduction of sleep duration following food intake at the beginning of their resting phase. Indeed, an acute 8-hour sleep deprivation has previously been shown to impair spatial memory consolidation in *M. murinus* (Rahman et al., 2013). This hypothesis is however unlikely to explain the present findings, as video recordings from the aviaries of the desynchronised animals showed that the entire food ration was consumed within 1 to 1.5 hours after food delivery, after which the animals returned to their nest boxes and slept. Hence, a reduction of only one to two hours of sleep per day is unlikely to induce significant deleterious effects in cognitive abilities in *M. murinus*. Some individuals also appear naturally more anxious or more sensitive to handling and experimental procedures than others, which may affect their performance in cognitive tasks. Increasing the sample size would therefore help reduce the influence of this inter-individual variability and improve the robustness of cognitive assessments.

### The potential role of metabolic flexibility in circadian resilience

Defined as an organism’s ability to adjust the oxidation of energy substrates to changing nutritional and energy conditions, metabolic flexibility enables the efficient transition from lipid utilisation during fasting to carbohydrate utilisation in the postprandial state (Goodpaster & Sparks, 2017). Thus, the metabolic adaptations observed in grey mouse lemurs should therefore be interpreted in the light of their evolutionary history. Unlike most models used in chronobiology, including laboratory rodents and humans, *M. murinus* is a nocturnal heterothermic primate that evolved under the highly challenging environmental conditions of the Madagascan forests, where ambient temperatures and food availability fluctuate markedly across seasons (Giroud et al., 2008; Schmid, 2000). These ecological constraints have likely favoured the evolution of exceptional metabolic flexibility, characterised by the capacity to efficiently alternate between energy storage, reserve mobilisation, and profound reductions in energy expenditure during torpor. Such metabolic hyper-flexibility enables grey mouse lemurs to continuously adjust substrate utilisation in response to environmental challenges, potentially protecting them against metabolic disorders associated with circadian misalignment. Conversely, impaired metabolic flexibility is commonly observed in obese and insulin-resistant humans and rodents, contributing to ectopic lipid accumulation, hepatic steatosis and dysregulation of glucose homeostasis (Goodpaster & Sparks, 2017; Koves et al., 2008). Reduced metabolic flexibility may therefore exacerbate the adverse effects of circadian disruption, particularly when food intake occurs at biologically inappropriate times. The grey mouse lemur could thus represent a valuable model for exploring the mechanisms underlying metabolic resilience to circadian-feeding misalignment, and for identifying factors that may contribute to inter-individual differences in vulnerability to metabolic disorders in humans.

Furthermore, as metabolic and behavioural adaptations such as torpor, huddling or the accumulation of fat reserves are strongly influenced by sex, age and season in *M. murinus* (Hozer et al., 2019; Noiret et al., 2020, 2024), it would be interesting to investigate the effects of circadian-feeding misalignment in males, during the winter season, or in older animals. Such studies would be particularly relevant given that ageing is associated with a progressive decline in physiological and metabolic flexibility in this species (Terrien et al., 2009). A valuable next step would be to investigate the effects of combined disruption of feeding and photic cues. As the light-dark cycle remained intact in the present study, central circadian synchronisation was likely preserved. Exposure to constant darkness or light could therefore reveal whether the metabolic and cognitive resilience observed in *M. murinus* depends on the maintenance of central clock entrainment.

## Conclusion

Overall, a 10-week period of circadian-feeding misalignment in adult female *Microcebus murinus* did not induce major alterations in physiological, metabolic or cognitive function. Although changes were observed in insulin, triglyceride and HDL levels, grey mouse lemurs appeared to adapt efficiently to the shifted feeding schedule, notably through a reversal in the temporal pattern of substrate utilisation, while maintaining their capacity to switch between energy sources. Because of the limited sample size and statistical power inherent to studies in non-human primates, subtle physiological or cognitive effects may have remained undetected. Nevertheless, these findings highlight the remarkable metabolic flexibility of this species and suggest that such flexibility may contribute to resilience against the adverse effects of circadian-feeding misalignment. In this context, elucidating the physiological and molecular mechanisms underlying this adaptive capacity represents an important challenge for future research on circadian desynchronisation and metabolic health.

## Acknowledgements

The expert animal care and management provided by A. Anzeraey, L. Dezaire, E. Pasquier, I. Hiron and J.M. Héliès is gratefully acknowledged. We also thank O. Le Carre and S. Pham for their helpful assistance during the study.

## Author contributions

Conceptualization: Jérémy Terrien, Océane Lavaux Investigation: Océane Lavaux, Eva Schneider, Coraline Ruiz Methodology: Jérémy Terrien, Océane Lavaux, Coraline Ruiz Resources: Océane Lavaux, Jérémy Terrien, Coraline Ruiz Supervision: Jérémy Terrien Writing – original draft: Océane Lavaux, Jérémy Terrien Writing – final review: Océane Lavaux, Jérémy Terrien, Coraline Ruiz, Eva Schneider

## Supplementary data

**Supp Figure 1:**
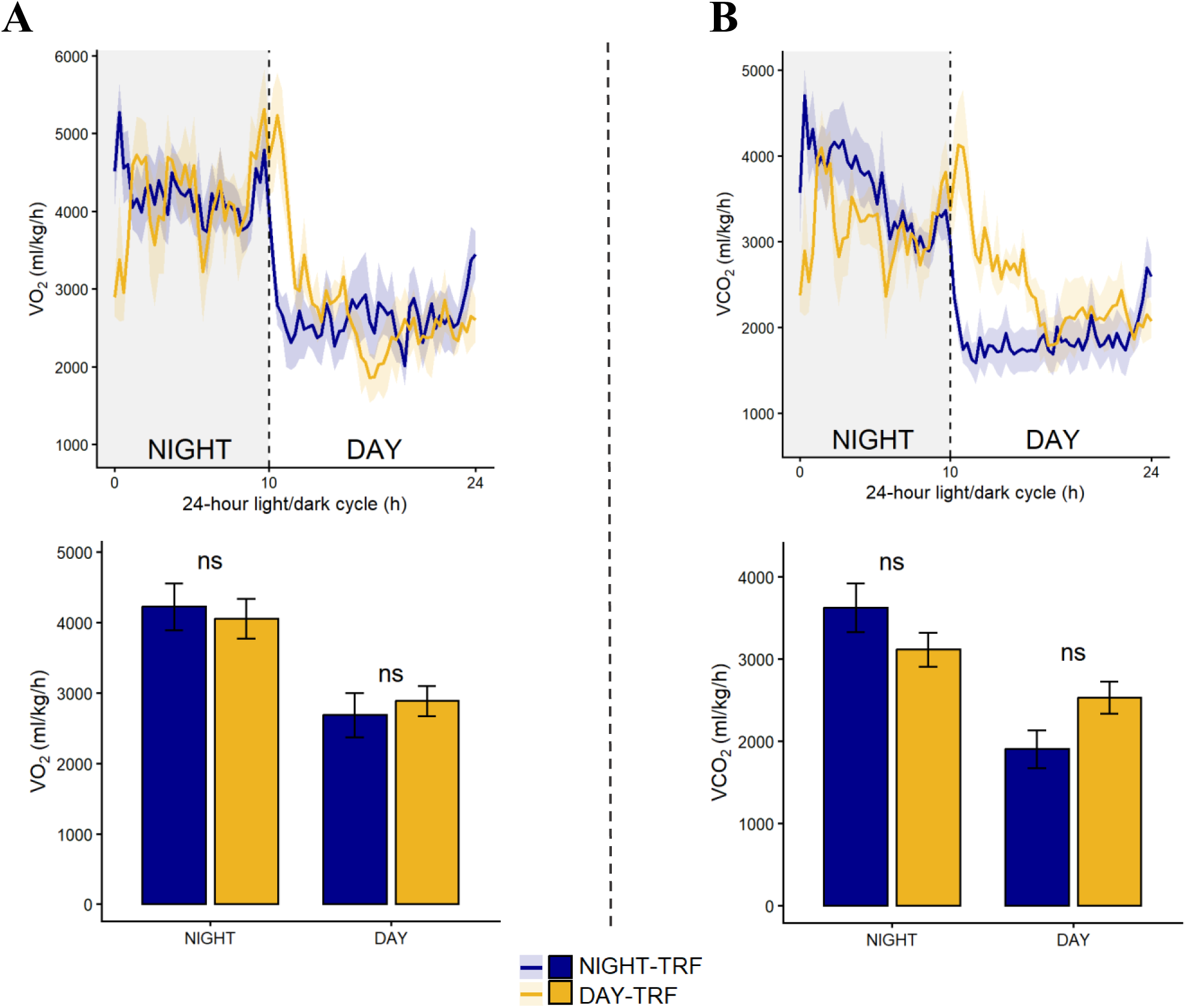
Complementary results of the indirect calorimetry assessment performed at week 9 of the study in adult female grey mouse lemurs, subjected or not to feeding-induced circadian desynchronisation. A/ VO_2_ (ml/kg/h): mean daily profiles (top) and averaged values (bottom) according to light phase (DAY VS NIGHT) and feeding schedule (NIGHT-TRF: n = 6; DAY-TRF: n = 5) after 9 weeks of treatment. Statistical results correspond to post-hoc comparisons between light phases and feeding schedule. B/ VCO_2_ (ml/kg/h): mean daily profiles (top) and averaged values (bottom) according to light phase (DAY VS NIGHT) and feeding schedule (NIGHT-TRF: n = 6; DAY-TRF: n = 5) after 9 weeks of treatment. Statistical results correspond to post-hoc comparisons between light phases and feeding schedule

**Supp Figure 2:**
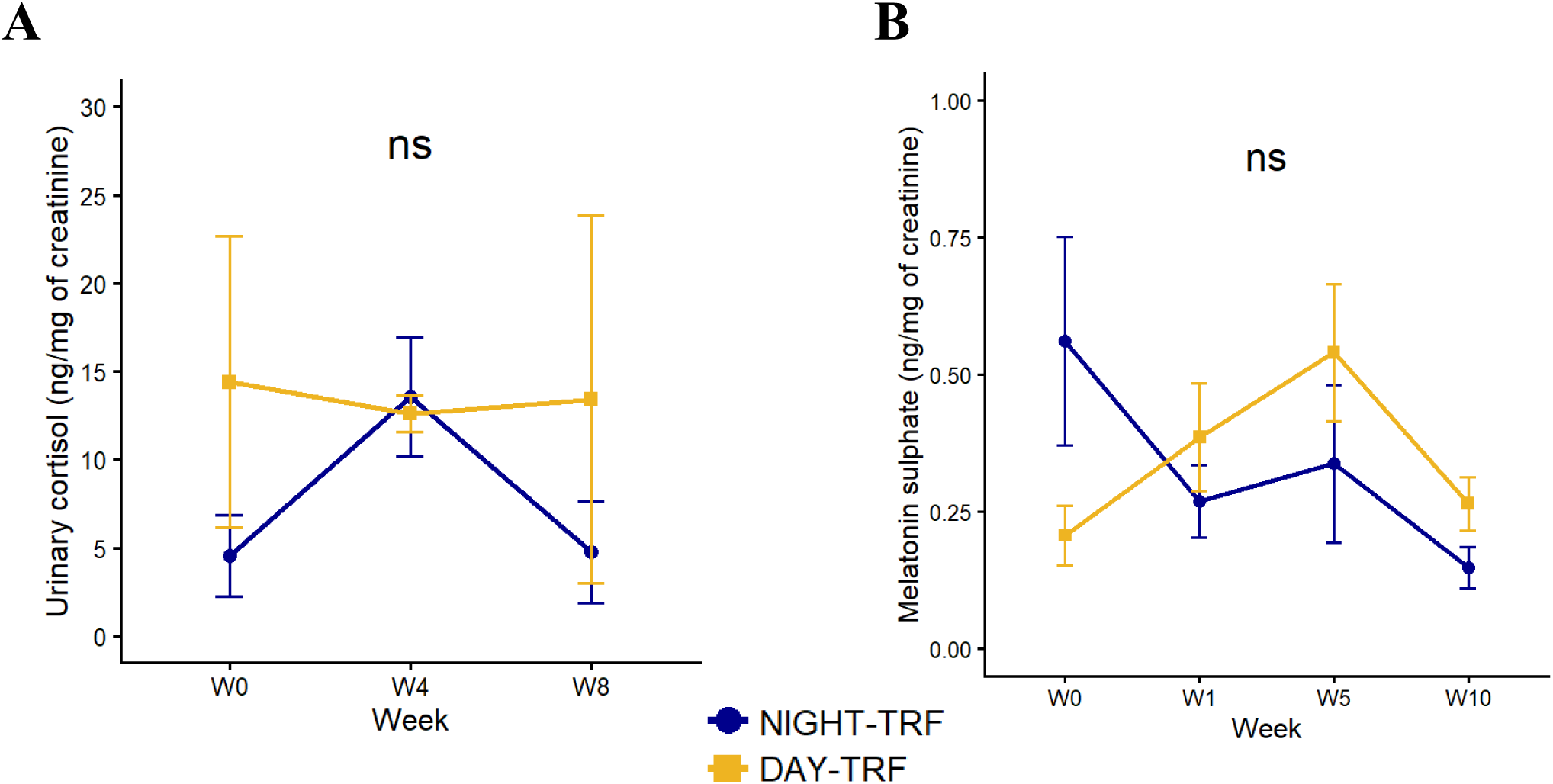
Complementary results of urinary cortisol and melatonin sulphate assays in adult female grey mouse lemurs subjected to circadian desynchronisation (DAY-TRF: n = 6) or maintained under a synchronised feeding schedule (NIGHT-TRF: n = 6). Both cortisol and melatonin sulphate values were log-transformed prior to analysis using linear mixed-effects models. A/ Changes in urinary cortisol concentrations (ng/mg of creatinine) over time (weeks). No significant effect of feeding schedule was detected (p = 0.43). B/ Changes in urinary melatonin sulphate concentrations (ng/mg of creatinine) over time (weeks). No significant effect of feeding schedule was detected (p = 0.42).

**Supp Figure 3:**
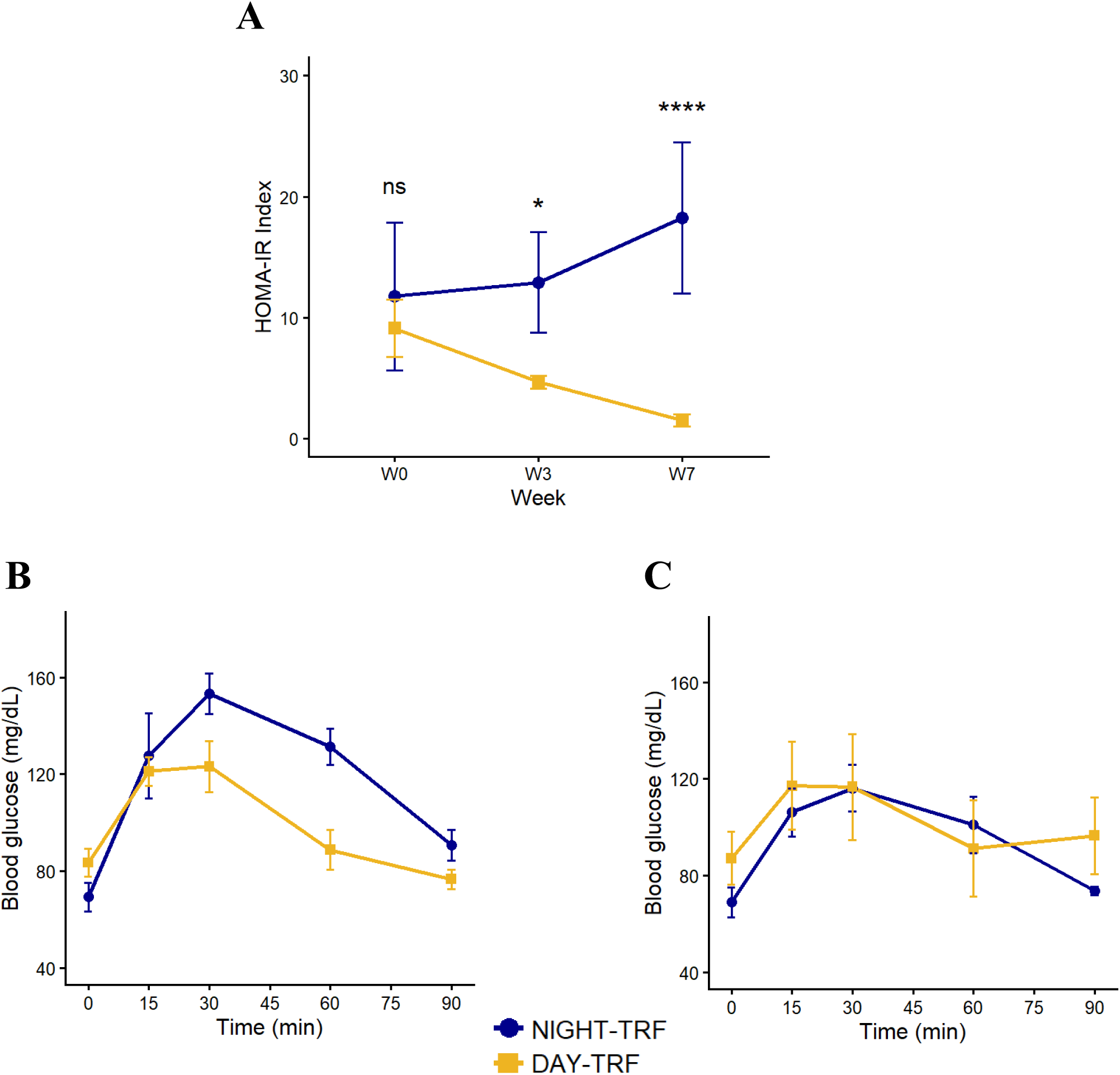
Complementary results regarding glucose homeostasis in adult female grey mouse lemurs subjected to circadian desynchronisation (DAY-TRF: n = 6) or maintained under a synchronised feeding schedule (NIGHT-TRF: n = 6). Data were analysed using linear mixed-effects models; HOMA-IR values were log-transformed prior to analysis. A/ Changes in HOMA-IR values over the course of the study. Statistics indicate the effect of feeding schedule within each week (post-hoc tests). B/ Mean blood glucose response curves during the oral glucose tolerance test (oGTT) performed at week 0 in both groups. C/ Mean blood glucose response curves during the oral glucose tolerance test (oGTT) performed at week 7 in both groups.

**Supp Figure 4:**
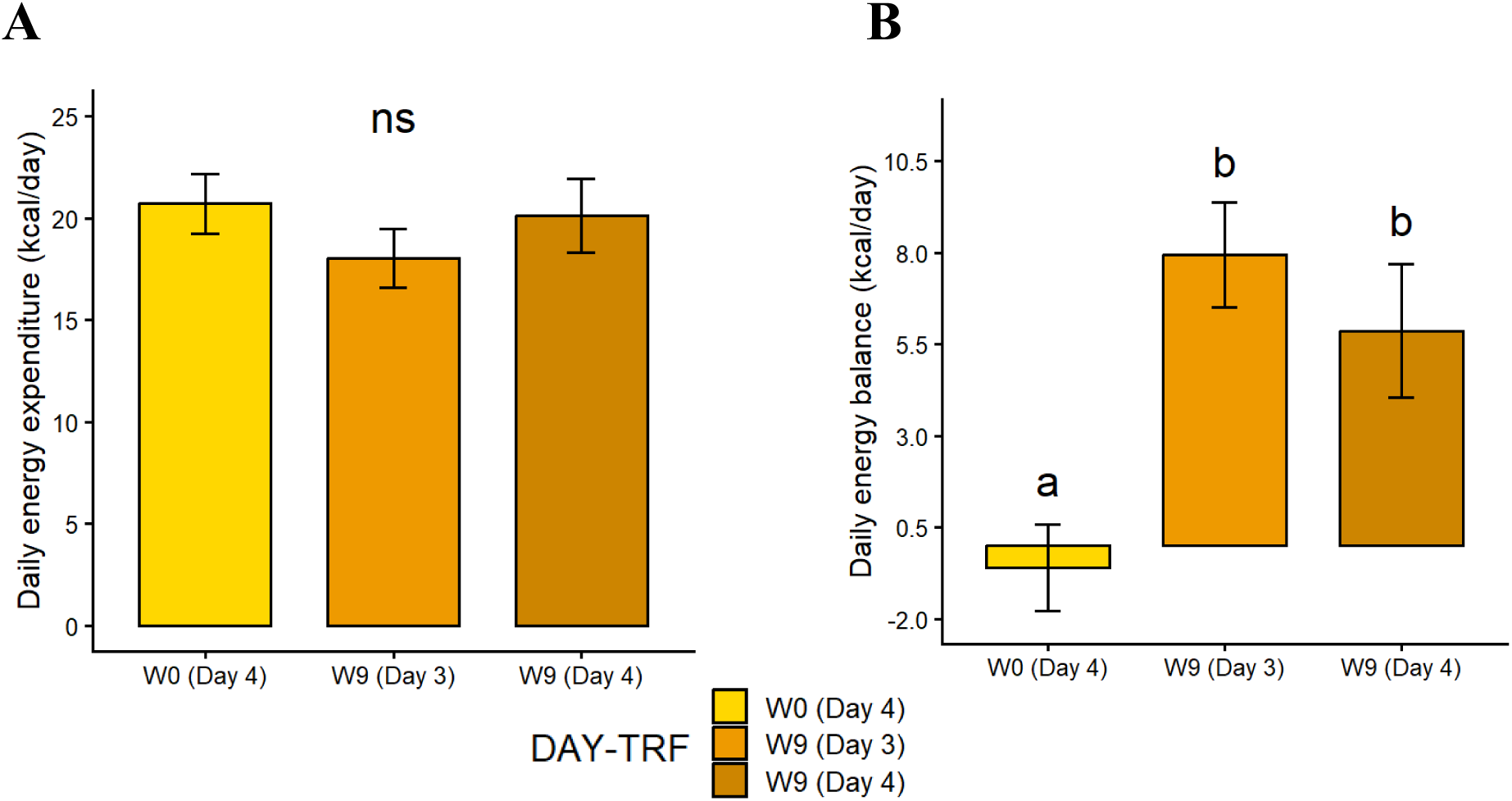
Complementary results regarding daily energy expenditure and balance within the DAY-TRF group where adult female grey mouse lemurs were subjected to circadian-feeding desynchronisation (W9 Day 3 and W9 Day 4) or not yet (W0 Day 4), analysed over different days of week 0 and week 9 indirect calorimetry sessions. A/ Daily energy expenditure (kcal/day). No statistical difference was detected between the different dates (p = 0.074). B/ Daily energy balance (kcal/day). Statistical results correspond to post-hoc comparisons between the dates: no difference was detected between W9 (Day 3) and W9 (Day 4) (p = 0.25) but both dates were statistically different from W0 (Day 4) (p < 0.001).

